# *Drosophila* sensory cilia lacking MKS-proteins exhibit striking defects during development but only subtle defects in adults

**DOI:** 10.1101/058081

**Authors:** Metta B. Pratt, Joshua S. Titlow, Ilan Davis, Amy R. Barker, Helen R. Dawe, Jordan W. Raff, Helio Roque

**Author notes:** Joint-Corresponding Authors.

## Abstract

Cilia are conserved organelles that have important motility, sensory and signalling roles. The transition zone (TZ) at the base of the cilium is critical for cilia function, and defects in several TZ proteins are associated with human congenital ciliopathies such as Nephronophthisis (NPHP) and Meckel Gruber syndrome (MKS). In several species, MKS and NPHP proteins form separate complexes that cooperate with Cep290 to assemble the TZ, but flies lack an obvious NPHP complex. We show that MKS proteins in flies are spatially separated from Cep290 at the TZ, and that flies mutant for individual *MKS* genes fail to recruit other MKS proteins to the TZ, while Cep290 appears to be recruited normally. Although there are abnormalities in microtubule and membrane organisation in developing *MKS* mutant cilia, these defects are less apparent in adults, where sensory cilia and sperm flagella function quite normally. Thus, surprisingly, MKS proteins are not essential for cilia or flagella function in flies.

## Introduction

Cilia and flagella are microtubule (MT)-based extensions of the plasma membrane present in evolution since the last eukaryotic common ancestor (LECA) (Ishikawa and Marshall, 2011; Nigg and Raff, 2009; Sung and Leroux, 2013). Cilia are present in many cell types and have diverse roles in cellular sensation, signalling and motility (Basten and Giles, 2013; Berbari et al., 2009; Nigg and Raff, 2009). Cilia are formed when the centriole pair migrates to the plasma membrane (PM). The older, mother, centriole forms a basal body (BB) that docks at the PM and MTs extend from the distal end of the BB to form a membrane-bounded axoneme. Concomitantly, many proteins are recruited to the Transition Zone (TZ), a complex structure often containing Y-links and transition fibres, that is assembled where the BB meets the PM (Garcia-Gonzalo and Reiter, 2012; Ishikawa and Marshall, 2011). The TZ is thought to be essential for cilium function and it appears to help assemble a membrane and cytoplasmic barrier that allows the cilium to form a distinct cellular compartment (Czarnecki and Shah, 2012; Hsiao et al., 2012; Hu and Nelson, 2011; Nachury et al., 2010; Reiter et al., 2012).

Interestingly, many of the genes linked to cilia dysfunction in humans, such as those associated with Meckel Gruber syndrome (MKS), Joubert Syndrome (JBT) and Nephronophthisis (NPHP), encode proteins that localise to the TZ (Czarnecki and Shah, 2012; Davis and Katsanis, 2012; Hu and Nelson, 2011; Reiter et al., 2012). Several studies have indicated that the TZ is formed by a large number of proteins that appear to function in broadly three distinct complexes or “modules”, an MKS module (comprising proteins such as MKS1, Tectonic, B9D1 and B9D2), an NPHP module (comprising proteins NPHP1 and NPHP2) and a Cep290 module (comprising proteins such as CEP290 and ATXN10)—although the precise components of each module may vary depending on cell type and species, and there may be some overlap of the proteins found within each module (Garcia-Gonzalo et al., 2011; Chih et al., 2012; Sang et al., 2012; Schouteden et al., 2015; Yee et al., 2015; Williams et al., 2008, 2011). Studies primarily in worms and mice suggest that proteins of one module are generally required for the recruitment of other members of the same module, while the recruitment of each module can occur at least partially independently of the other modules (Basiri et al., 2014; Schouteden et al., 2015; Williams et al., 2008, 2011). Moreover, studies in worms indicate that there is a modular hierarchy of TZ assembly, with the Cep290 module at the top and the MKS and NPHP modules below. Mutating or depleting components from either the MKS or the NPHP module have a relatively mild effect on cilia function, while co-mutating or co-depleting components of both modules at the same time has a much stronger effect (Schouteden et al., 2015; Williams et al., 2008).

A recent genomic analysis of the evolutionary conservation of TZ components suggested that just 6-7 components of the MKS module (TMEM67, CC2D2A, B9D1, B9D2, AHI1, Tectonic and MKS1) comprise a core conserved module for TZ assembly, although even these 6-7 proteins do not appear to be present in every species that forms a cilium with a TZ. Thus, many questions remain as to how the proteins of the TZ work together to form a functional TZ and how this process goes awry in human ciliopathies.

The fruit fly *Drosophila melanogaster* has proved a powerful experimental system for many years, yet comparatively little research has focused on the TZ in flies. The fly offers several advantages as an experimental system to study TZ function. Perhaps most importantly, flies do not use cilia for hedgehog or wingless signalling, so flies lacking cilia develop largely normally, without the gross morphological perturbations associated with defects in these signalling pathways in vertebrate systems. Indeed, most adult fly cells do not contain cilia/flagella, which are restricted to certain sensory neurons and sperm lineages. Adult flies lacking centrioles and cilia are largely morphologically normal, although they are severely uncoordinated due to the lack of cilia in their mechano-sensory neurons and they die shortly after eclosure. Moreover, while several TZ proteins have been identified in flies, including Cep290 and several members of the MKS module (including all but one of the “core” MKS module proteins: TMEM67, CC2D2A, B9D1, B9D2, Tectonic and MKS1—but not AHI1), few, if any, NPHP module proteins have been identified, suggesting that flies may rely only on the CEP290 and MKS1 modules for TZ assembly, potentially simplifying the analysis of TZ assembly and function (Barker et al., 2014; Basiri et al., 2014).

It has recently been shown that Cep290 and Chibby (a conserved TZ protein that is also involved in Wnt signalling in vertebrates, but not in flies) are required for cilia function in flies (Basiri et al., 2014; Enjolras et al., 2012). Cep290 and Cby mutants are uncoordinated and have reduced male fertility and they exhibit structural defects in their sensory neuron cilia and in the short primary cilia found in maturing spermatocytes as well as in the axonemes of mature sperm. In Cep290 mutants several MKS module proteins (MKS-1, B9D1 and B9D2) were still recruited to the growing ciliary cap structure in elongating spermatids, although their localisation was abnormally diffuse. The function of the conserved MKS module of proteins, however, has not been directly addressed in flies.

Here, we analyse the distribution of several MKS-module proteins in flies and generate mutations in two of the genes encoding these proteins, *MKS1* and *B9D1.* We show that the MKS-proteins are recruited to an area of the TZ that is spatially distinct from Cep290, and that mutations in *MKS1* and *B9D1* disrupt the TZ localisation of the other MKS-proteins, supporting the idea that these proteins form a functional complex. Despite the lack of detectable MKS proteins at the TZ, *MKS1* and *B9D1* mutants are viable and fertile and, although MKS1 mutants exhibit clear structural defects at sensory cilia during earlier stages of development, these defects are largely absent in adults. We conclude that even though flies lack an obvious NPHP module, they can still form functional cilia and flagella without an MKS module, given enough developmental time.

## Results

### TZ proteins in *Drosophila melanogaster*

We previously performed a bioinformatics analysis of putative TZ proteins across species and identified a core conserved group of proteins that are present in >50% of ciliated organisms: AHI1, B9D1, B9D2, CC2D2A, Tectonic, TMEM67 and likely MKS1 (Barker et al., 2014). *Drosophila melanogaster* has identifiable homologues of all of these proteins except AHI1, and also has homologues of the TZ proteins Cep290, TMEM216, TMEM231 and TMEM237; no components of the core NPHP TZ module were identified (Barker et al., 2014). These findings are in broad agreement with another bioinformatics analysis of TZ proteins in *Drosophila* (Basiri et al., 2014). In addition, Chibby (Cby) and Dilatory (Dila) have also recently been identified as components of the *Drosophila* TZ (Enjolras et al., 2012; Ma and Jarman, 2011).

### *Drosophila* TZ proteins occupy distinct regions in spermatocyte cilia

To better understand how TZ proteins are organized in *Drosophila* cilia we generated fly lines individually expressing GFP-fusions to the core MKS-module proteins MKS1, B9D1, B9D2, TMEM216, CC2D2A and Tectonic. Protein expression was driven from the ubiquitin promoter that is expressed at moderate levels in all fly tissues (Lee et al., 1988) with the exception of CC2D2A which was driven from its endogenous promoter. We also obtained lines expressing the TZ proteins Cep290-GFP (Basiri et al., 2014) and Chibby (Cby)-GFP (Enjolras et al., 2012). We used dual colour 3D super resolution structured illumination microscopy (3D-SIM) to image the distribution of each GFP-fusion at the BB/axoneme *in vivo* in fixed mature spermatocytes labelled with anti-Asterless (Asl) antibodies to mark the outer wall of the BB (Franz et al., 2013; Varmark et al., 2007) (Figure 1A). These spermatocytes have very long BBs that extend a short axoneme (Tates, 1971) (Figure S1), and several genes encoding TZ proteins are highly expressed in testes. Moreover, mutations in Cep290, Cby and Dila all lead to defects in spermatogenesis (Basiri et al., 2014; Enjolras et al., 2012; Ma and Jarman, 2011).

**Figure 1.**
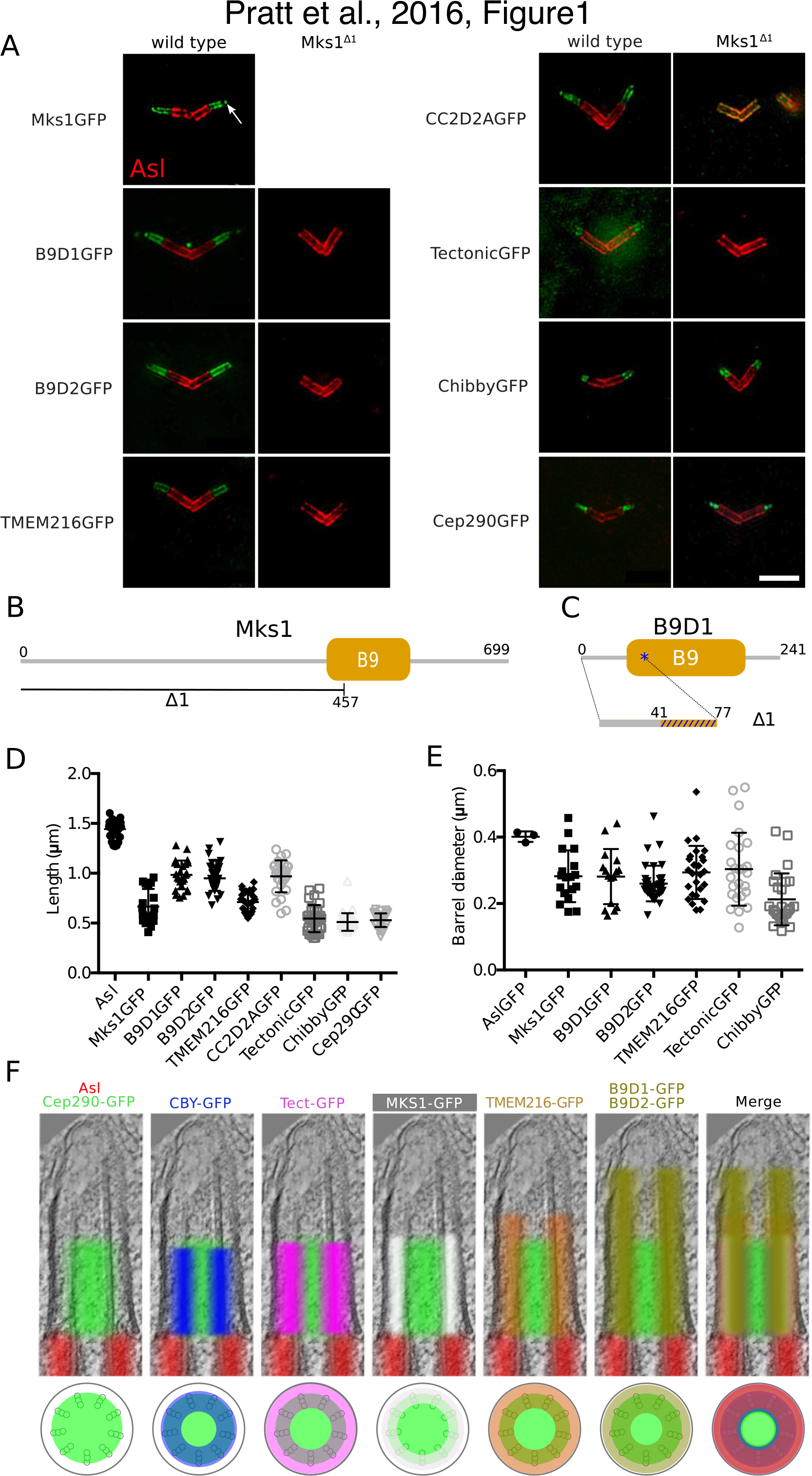
MKS proteins occupy distinct regions within the TZ and depend on MKS1 for their T3Z localization. (**A**) 3D-SIM micrographs of *Drosophila* spermatocyte cilia showing the localisation of various GFP-tagged TZ proteins (*green,* as indicated) in relation to the BB protein Asterless (Asl) (*red*) in WT (left panels) and *Mks1^Δ1^* mutants (right panels). (**B,C**) Schematic representations of the *Mks1^Δ1^* deletion (B) and the *B9D1^Δ1^* mutation (C)—that creates a frameshift at amino acid 41 leading to a scrambled aa code and a premature stop codon at amino acid 77 (blue asterisk). (**D**) Graph quantifies the longitudinal length of Asl and the various TZ proteins in 3D-SIM micrographs. The isolated GFP dots, sometimes present at the distal tips of the TZ (arrows, in [A]), were ignored for these measurements. (**E**) Graph quantifies the diameter of the “barrel’ formed by Asl and the various TZ proteins in 3D-SIM micrographs (see Materials and methods). (**F**) Using the mean length and diameter measurements obtained in (D,E) we created composite images (top panels) superimposing the localisation of Asl (*red*) and the TZ proteins (*various colours,* as indicated) on a typical *Drosophila* spermatocyte EM micrograph. A schematic of the transverse profile of the BB/axoneme (lower panel) shows the various protein localizations in relation to the nine fold MT blades (grey triplets) and the axoneme membrane (black circle). Error bars represent the s.e.m. Scale bar=2 μjm.

In WT spermatocytes all of the TZ proteins were located distally to the BB (stained by Asl, *red,* Figure 1A) and extended into the axoneme to varying extents (Figure 1A). We measured the average length and width of these axoneme extensions (note that for CC2D2A-GFP we could not accurately measure the width as its distribution was too irregular) (Figure 1D,E). We then created compound images (Figure 1F) by overlaying the average distribution of each protein on an EM-tomogram of a typical wild-type (WT) BB/axoneme, aligning the markers so that the Asl staining terminated at the distal end of the BB (see Materials and Methods). These studies revealed that Cep290 occupied a distinct inner region of the TZ (shown in *green* in all the compound images) that appeared to overlap with the axonemal MTs. The distribution of Cby (*blue*) overlapped the outer portion of the Cep290 region, while Tectonic (*purple*) and MKS1 (*white*) occupied a region between the axonemal MTs and the ciliary membrane that was largely outside of the Cep290 region. TMEM216 (*orange*) and B9D1/B9D2 (*dark green*) all also occupied a similar region between the axoneme and the membrane, but these regions extended distally beyond the other TZ proteins, with B9D1 and B9D2 extending the furthest. These studies demonstrate that individual TZ proteins occupy distinct regions within the TZ, in agreement with recent findings in cultured RPE-1 (Tony Yang et al., 2015).

### MKS1 and B9D1 are required to localise the MKS module to the TZ

To study the function of the *Drosophila* MKS complex in TZ formation we generated an *Mks1* mutation by imprecise P-element excision. This generated a 1.4Kb deletion that removed the N-terminal 474aa of the MKS1 protein, including the start codon and part of the conserved B9 domain (Figure 1B), a domain of unknown function that is often found in cilia/flagella associated proteins. We hereafter refer to flies homozygous for this mutation as *Mks1^Δ1^* mutants.

We analysed the localization of GFP-fusions to the other TZ proteins by 3D-SIM in an *Mks1^Δ1^* background. The TZ localisation of Cep290-GFP and Cby-GFP was not detectably perturbed, but B9D1-GFP, B9D2-GFP, TMEM216-GFP and Tectonic-GFP were no longer detectable at the TZ in *Mks1^Δ1^* mutants (Figure 1A). Interestingly, although CC2D2A-GFP was also no longer detectable at the TZ in *Mks1^Δ1^* mutants, this protein appeared to re-localise to the BB walls (Figure 1A). These findings strongly suggest that the recruitment of the entire MKS-module of proteins to the TZ is dependent on MKS1, and that although a TZ that can recruit Cep290 and Cby is still formed in *Mks1^Δ1^* mutants, this TZ lacks detectable MKS-module proteins.

We also generated a mutation in the *B9D1* gene using CRISPR/Cas9 technology. This mutation generated a frame shift leading to the introduction of a premature stop-codon and so presumably to the effective deletion of the C-terminal 164aa of the B9D1 protein (Figure 1C); we hereafter refer to flies homozygous for this mutation as *B9D1^Δ1^* mutants. The TZ localisation of MKS1-GFP and B9D2-GFP were no longer detectable in *B9D1^Δ1^* mutants (Figure S2). As MKS1 is required for the localisation of all the other MKS-module proteins we have examined to the TZ, the absence of MKS1 from the TZ of *B9D1^Δ1^* mutants means that the other MKS module proteins are also likely absent from the TZ in *B9D1^Δ1^* mutants. Thus, in agreement with previous reports (Bialas et al., 2009; Williams et al., 2008), MKS-module proteins appear to be interdependent for their localization to the TZ, and MKS-module proteins are not detectable at the TZ in *Mks1^Δ1^* and *B9D1^Δ1^* mutants.

### <u>*Mks1^Δ1^* and *B9D1^Δ1^* mutants are viable, fertile and do not exhibit dramatic</u> <u>defects in sensory cilia function</u>

To our surprise, *Mks1^Δ1^* and *B9D1^Δ1^* mutants were not noticeably uncoordinated and we have maintained homozygous stocks of these mutants in the laboratory for more than two years, indicating that they are both male and female fertile. This is in contrast to previously described mutations in the TZ protein encoding genes *Cep290* (Basiri et al., 2014), *Cby* (Enjolras et al., 2012) and *dila* (Ma and Jarman, 2011) that are all severely uncoordinated due to defects in their sensory cilia, and which all exhibit severely reduced male fertility. In quantitative fertility tests *Mks1^Δ1^* mutant males were not significantly less fertile than WT controls, indicating that mutant sperm flagella were largely functional (Figure 2A).

**Figure 2.**
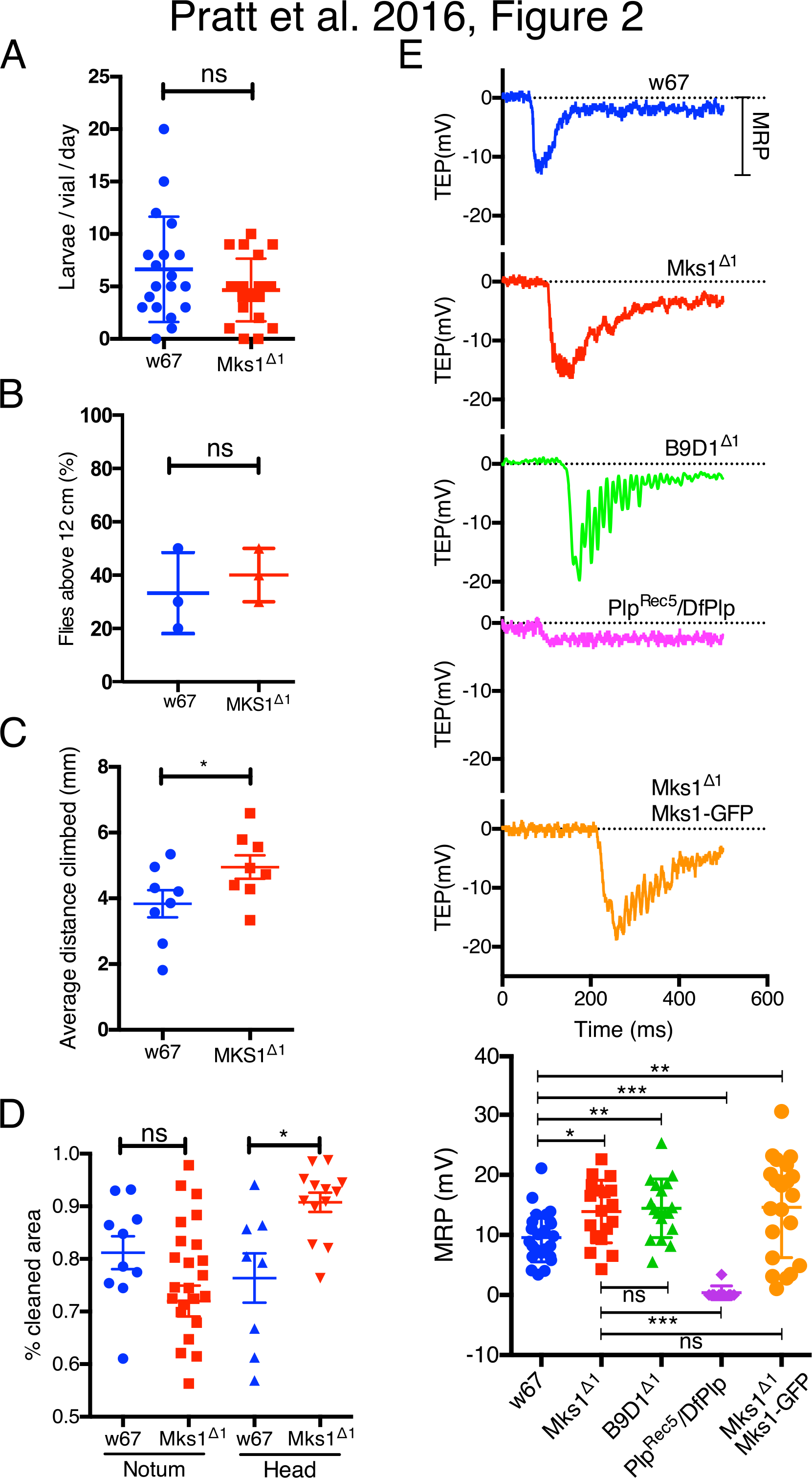
MKS-Module mutants exhibit no dramatic behavioural defects. (**A**) Graph quantifies a male fertility test assessing the number of larvae that hatched per vial per day when WT (*w^67^*) or *Mks1^Δ1^* mutant males were mated to WT females. (**B, C**) Graphs quantify gravitaxis tests assessing the climbing reflex of WT (w^67^) or *Mks1^Δ1^* mutant flies to climb to the top of a vial (B) and the speed at which flies climb (C). (D) Graph quantifies a cleaning reflex test assessing the ability of WT (w^67^) or *Mks1^Δ1^* mutant flies to clean dust from their notum or head. (E) Electrophysiology traces show typical examples of the trans-epithelial electron potential (TEP) of individual mechano-sensory bristles of the sensory notum cilia recorded from either WT, *Mks1^Δ1^, B9D1^Δ1^* or (as a negative control) *Plp^Rec5/^DfPlp* flies. A trace from an *Mks1^Δ1^* mutant rescued by the MKS1-GFP transgene is also shown. The graph (bottom panel) quantifies the average mechanical response potential (MRP) derived from these traces for each genotype. Error bars represent the s.d.; significance was assessed using a one-way ANOVA: ^*^p<0.01, ^**^p<0.001, ***p<0.0001.

We next performed a series of assays designed to test sensory cilia function. We first tested the flies geotaxis response (Hirsch and Tryon, 1956). When WT flies are knocked down to the bottom of a tube they quickly climb to the top, while flies with defective sensory cilia function are uncoordinated and cannot climb efficiently. There was no significant difference in the number of flies counted at the top of the vial 10secs after knock down between WT and *Mks1^Δ1^* mutants (Figure 2B), but we noticed that *Mks1^Δ1^* flies appeared to climb faster than WT. We therefore repeated the assay, but measured the average distance climbed per vial after 7s of knockdown. This confirmed that the *Mks1^Δ1^* mutants climb slightly faster than WT (Figure 2C). We next compared the dust grooming response of WT and *Mks1^Δ1^* mutant flies, which is driven by the mechanical stimulation of external bristles by dust (Phillis et al., 1993). This assay measures the cleaned area of the head and the notum, 30 and 90mins respectively, after applying dust to the flies. This revealed that *Mks1^Δ1^* flies had a significantly higher cleaned area on their heads but no difference was observed in the notum (Figure 2D). Taken together, these observations confirm that sensory cilia function in *Mks1^Δ1^* mutants is not dramatically perturbed and that, if anything, mutant flies exhibit a slightly increased response to certain stimuli (see below), although this could be due to genetic background differences between strains.

### <u>Mechanoreceptor potentials of *Mks1^Δ1^* and *B9D1^Δ1^* mutant sensory cilia</u> <u>are not dramatically perturbed</u>

To more directly examine sensory cilium function in *Mks1^Δ1^* mutants we measured the electrophysiological response of mechano-sensory bristles in the notum. Each of these bristles is innervated by a peripheral sensory neuron that extends a cilium at the base of the bristle. The supporting cells that encapsulate the cilium generate a trans-epithelial electron potential (TEP) that can be detected by placing one electrode over the cut end of a bristle and inserting a reference electrode into the thorax (Kernan, 1994). Deflecting the bristle with a mechanical stimulus causes ion channels in the cilium membrane to open, allowing ions to flow from the support cells and generate a mechanical response potential (MRP). MRPs in *Mks1^Δ1^* mutants and in *B9D1^Δ1^* mutants did not differ dramatically from controls, in fact both mutants exhibited a slight, but significant, increase in MRP amplitude (Figure 2E). As a negative control we made similar recordings in *Pericentrin-Like-Protein (PLP*) mutants that have previously been shown to have severely reduced sensory cilia function (Martinez-Campos et al., 2004)(Figure 2E). We also analysed a line expressing Mks1-GFP in the *Mks1^Δ1^* mutant background and observed that this strain also had a slight increase in the MRP response (Figure 2E), suggesting that the slight increase in sensitivity is either due to background differences between the WT and mutant strains analysed, or is not efficiently rescued by the Mks1-GFP transgene. In either case, it is clear that the electrophysiological response of these sensory cilia falls within the normal range (Dubruille, 2002) in *Mks1^Δ1^* and *B9D1^Δ1^* mutants.

### <u>*Mks1^Δ1^* mutant spermatocyte axonemes are shortened, but mutant</u> <u>testes exhibit no dramatic defects</u>

To better understand why the lack of MKS-module proteins appears to have only a very mild effect on cilia and flagella function we examined cilia and flagella structure by EM tomography. We first examined testes, as MKS module proteins are most highly expressed in this tissue and mutations in the TZ proteins Cep290, Cby and Dila all exhibit severe defects in testes (Basiri et al., 2014; Enjolras et al., 2012; Ma and Jarman, 2011). An EM tomography examination of *Mks1^Δ1^* mutant flagella revealed no obvious structural defects, consistent with our observations that mutant flies are male fertile (Figure S3). However, we noticed that the short axonemes that are normally assembled in mature spermatocytes (Tates, 1971) were dramatically shortened in mutant cells (Figure 3A,C,D, Movie S1,S2), while the length of the very long BBs at the base of these axonemes was not dramatically affected (Figure 3B). Importantly, this short axoneme phenotype was rescued by the expression of the Mks1-GFP transgene (Figure 3A,C,D, Movie S3). These findings strongly suggest that the proper assembly of spermatocyte axoneme is dependent on MKS-module proteins.

**Figure 3.**
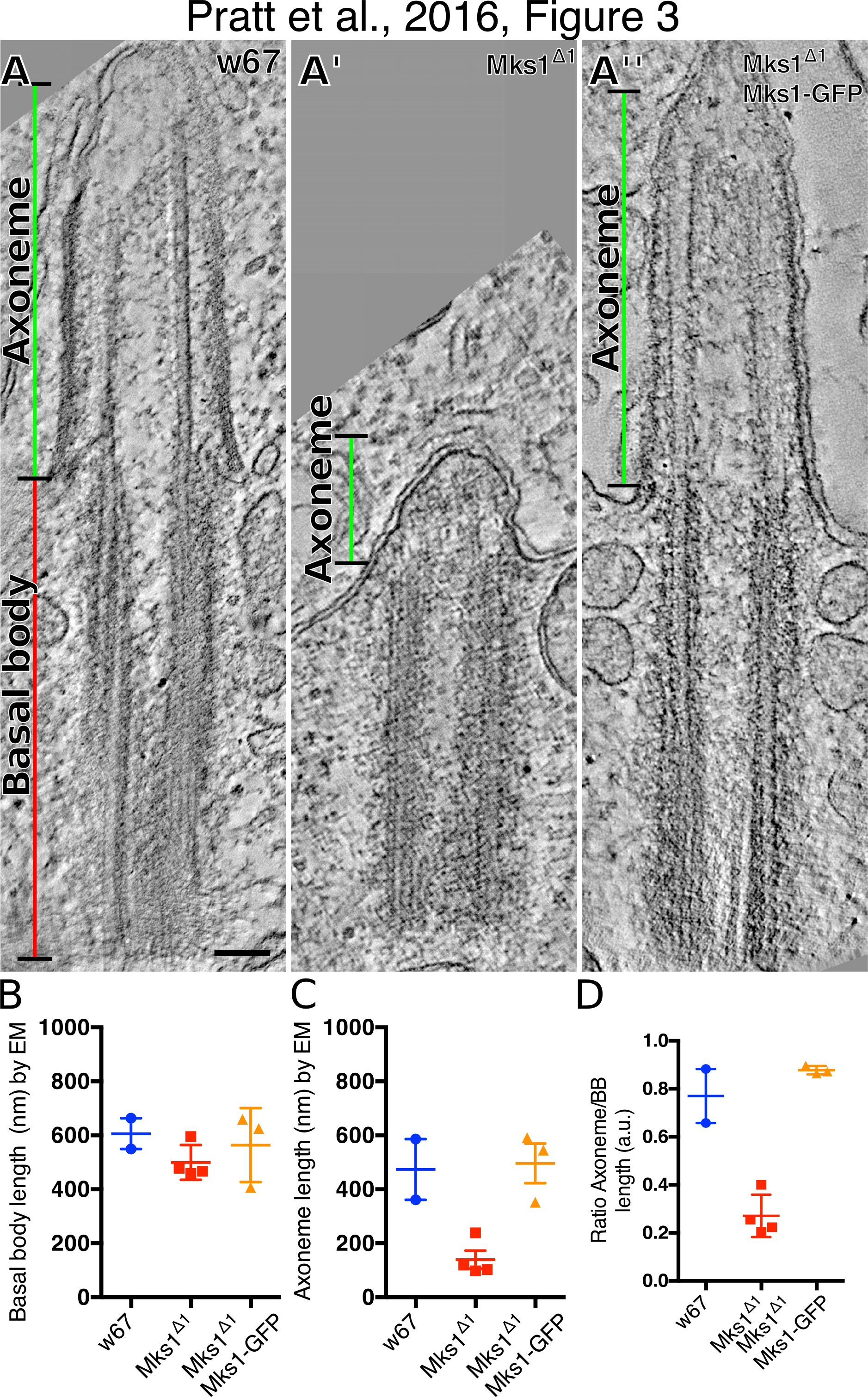
*Mks1^Δ1^ mutant primary spermatocytes have short axonemes.* (**A-A’’**) Micrographs show electron tomograms of primary spermatocyte cilia from WT (A), *Mks1^Δ1^* mutant (A’) and *Mks1^Δ1^* mutant rescued by Mks1-GFP (A’’). (**B-D**) Graphs show the quantification of basal body (BB) length (B), axoneme length (C) and the ratio of the axoneme/BB length in WT (*blue* dots), *Mks1^Δ1^* mutant (*red* dots) and *Mks1^Δ1^* mutant rescued by MKS1-GFP (*orange* dots) measured from EM tomograms. Error bars represent the s.d.; note that only 2-4 tomograms from each genotype contained the entire BB/Axoneme complex, allowing us to make these measurements, so we have performed no statistical analysis of significance on this small number of samples. Scale bar=100nm.

### <u>The localisation of the trans-membrane proteins NompA and NompC is</u> <u>not detectably perturbed in *Mks1^Δ1^* mutant sensory cilia</u>

The slight increase in signalling response in the bristle hairs suggested a possible defect in the ion channel composition in the ciliary membranes of *Mks1^Δ1^* mutants. We therefore examined the distribution of the transmembrane proteins NompA and NompC. NompA is localised to the dendritic cap at the distal tip of the cilium and is essential for cilia function (Chung et al., 2001; Kernan, 1994) while NompC forms ion channels in the ciliary membrane that are enriched towards the distal tip of the cilium (Lee et al., 2010; Zhang et al., 2013). We looked at the notum of flies 72h after pupae formation (APF) and found that in both WT and *Mks1^Δ1^* mutant cells NompA-GFP localised strongly to the base of the bristle, with a dimmer and elongated dot in the bristle hair (Figure 4A, B), while NompC-GFP localized to a single dot at the bristle base (Figure 4C). This suggests that the distribution of ion channels in the ciliary membrane is not dramatically perturbed in *Mks1^Δ1^* mutant cilia.

**Figure 4.**
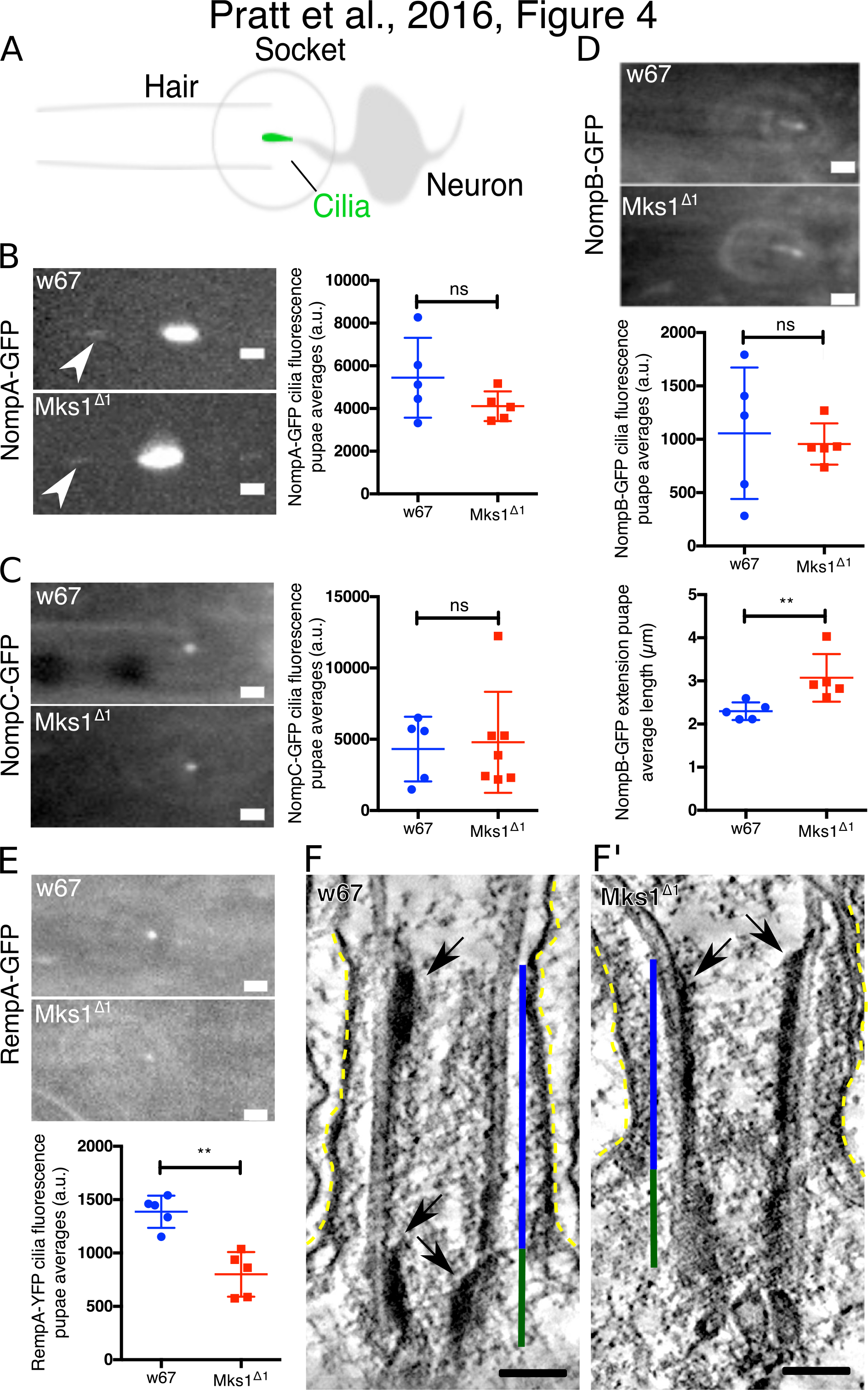
Mks1^Δ1^ mutant sensory cilia appear to have subtle defects in the localisation of IFT proteins. (**A**) A schematic illustration of the *Drosophila* notum hair socket and associated sensory cilia. (**B-E**) Micrographs show images and graphs show quantification of fluorescence intensity of the transmembrane proteins NompA-GFP (B) (which localises to the dendritic cap of the cilium), NompC-GFP (C) (a ciliary ion channel protein), RempA-GFP (the *Drosophila* homolog of IFT144) and NompB-GFP (the *Drosophila* homolog of IFT88) in WT and *Mks1^Δ1^* mutant sensory cilia. The localisation of NompA-GFP and NompC-GFP was not detectably perturbed in *Mks1^Δ1^* mutants, but significantly less RempA-GFP was localised in the *Mks1^Δ1^* mutant cilia, while an analysis of the average length of NompB-GFP staining (E, bottom panel) revealed that this was slightly longer in *Mks1^Δ1^* mutants. (**F,F’**) Micrographs show electron tomograms of the BB (*dark green* lines) and TZ (*blue* lines) in a WT (F) and an *Mks1^Δ1^* mutant (F’) sensory cilium (ciliary membrane is highlighted by *dotted yellow* lines). Electron dense particles similar to IFT trains (arrows) are visible on the BB microtubules facing the TZ lumen in WT. In *Mks1^Δ1^* sensory cilia these particles appear to be dramatically extended. Error bars represent s.d.; significance was assessed using a two-tailed t-test. Scale bars=2μm for B-D and100nm for F and F’.

### <u>Intraflagellar transport appears to be subtly perturbed in *Mks1^Δ1^* mutant</u> <u>cilia</u>

Our results so far suggest that sensory cilia lacking MKS-module proteins exhibit only mild functional defects. Previous studies suggested that MKS proteins interact with intraflagellar transport (IFT) proteins in zebrafish cilia (Zhao and Malicki, 2011). To test this possibility more directly in flies, we examined the localization of NompB-GFP—the IFT88 homolog responsible for anterograde transport (Han et al., 2003)—and of RempA-YFP—the IFT140 homolog that is part of the retrograde transport system (Lee et al., 2008a)—at the base of the notum bristle. In WT cells, NompB-GFP localized as a line at the base of the cilium that was enriched at the proximal end of the axoneme, and this was also the case in *Mks1^Δ1^* mutants, although the NompB-GFP signal was slightly, but significantly, extended (Figure 4D). RempA-YFP localized as a dot at the base of the bristle hair in both WT and mutant cells, although its intensity was significantly reduced in mutants (Figure 4E). These studies support the possibility that there are subtle defects in IFT in *Mks1^Δ1^* mutant cells.

IFT particles are often visible by EM as electron dense “IFT trains” located between the MTs and the ciliary membrane (Rogowski et al., 2013; Stepanek and Pigino, 2016; Vannuccini et al., 2016). Interestingly, in our EM analysis of WT cells we could not find such particles localized between the B-MTs and the ciliary membrane in the axoneme, but we instead found similar extended electron dense particles localizing along the A-MTs and facing the internal lumen of the BB/TZ (arrows, Figure 4F). These findings raise the intriguing possibility that IFT particles may assemble on the intraluminal surface of the BB in these cells. Supporting the idea that these particles might be IFT particles, we observed similar particles in *Mks1^Δ1^* mutant at 72h APF (Figure 4F) but these particles appeared to accumulate at the BB, forming a continuous sheet from the proximal lumen of the BB and into the TZ (137±32.5nm in 9 particles out of 3 cilia for WT vs. 184.8±70.1nm in 11 particles out of 3 cilia for Mks1^Δ1^)—in accordance with the increase in length of the NompB-GFP signal in *Mks1^Δ1^* mutant sensory cilia (Figure 4D). Together these results suggest that IFT is subtly perturbed in *Mks1^Δ1^* mutant sensory cilia.

### <u>*Mks1^Δ1^* mutant cilia exhibit striking structural defects at 48h APF, but are</u> <u>largely normal by 72h APF</u>

We next used ET to compare the ultrastructure of the sensory cilia in the bristles of the notum in WT and *Mks1^Δ1^* mutants at 48h and 72h APF (Figure 5). At 48h APF WT cilia (n=3) form relatively straight cilia that extend towards the plasma membrane of the hair cell (Figure 5A,B, Movie S4). At their base, these cilia have a basal body (BB—MTs of BB shown in *dark green*) that extends both A and B MT doublets into the TZ (*blue*) and then into the ciliary membrane (*light green*). The TZ is discernible as a straightening (*black* bar, Figure 5C) and electron-dense thickening (*blue* bar, Figure 5C) of the membrane, similar to that visualized in spermatocyte cilia (Figure 3A). The ciliary MTs doublets extend for ~2/3 of the cilia length and then continue as singlets of the A MTs; most of these A MTs terminate close to the tip of the cilium. Some MTs that were not connected to the BB were also found in the axoneme in close apposition to the ciliary membrane (*purple*) and an electron-dense crystalline structure surrounded the top ~1/3 of the axoneme, apparently linking the outer ciliary membrane with the adjacent sheath cell membrane (arrows, Figure 5D). At the base of the cilium, a daughter centriole was positioned just below the BB and it was surrounded by ciliary rootlets (Chen et al., 2015; Styczynska-Soczka and Jarman, 2015) emanating from the distal end of the BB (Figure 5D) where vesicles could also be observed (asterisks, Figure 5E).

**Figure 5.**
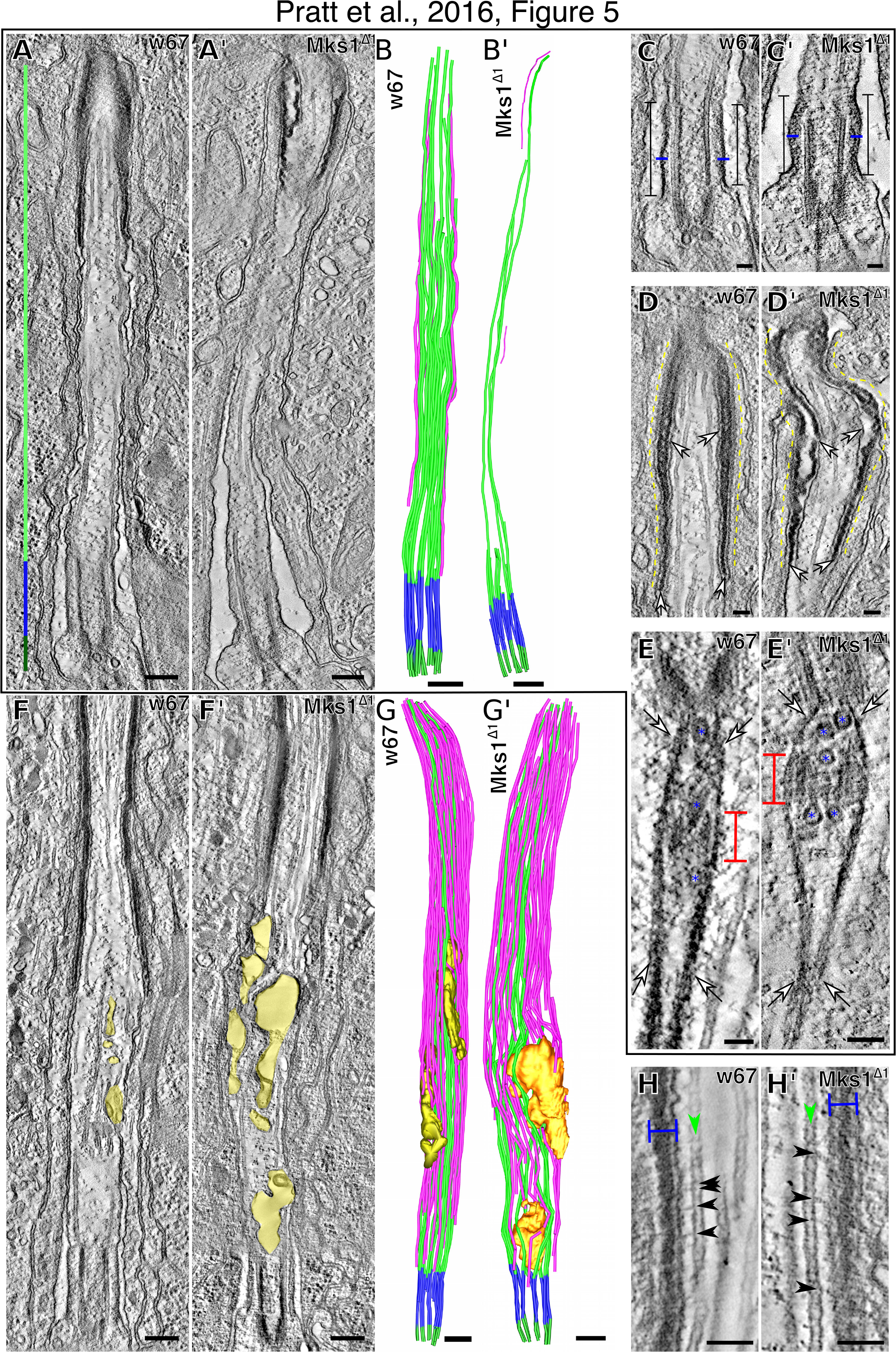
*Mks1^Δ1^* mutant sensory cilia exhibit structural defects at 48h APF, but are relatively normal by 72h APF. (**A,A’**) Micrographs show electron tomograms of the sensory cilium at 48h APF in wild type (A) and *Mks1^Δ1^* mutants (A’). (**B,B’**) Illustrations show the corresponding 3D models of these sensory cilia, highlighting the BB MTs (*dark green*), the transition zone MTs (*blue*) and axonemal MTs (*light green*). Axoneme MTs not directly attached to the BB are highlighted in *magenta*. (**C,C’**) Higher magnification electron tomograms of the TZ (square capped lines) in WT (C) and *Mks1^Δ1^* mutants (C’); the TZ is shorter in the mutant and the electron dense TZ membrane (*blue* lines) appears wider and more diffuse. (**D,D’**) At the tip of the WT axoneme there is a crystalline-like electron dense structure (E) (arrows) located between the ciliary membrane (*black* arrowheads) and the sheath cell membrane (*white* arrowheads); this structure is present in the *Mks1^Δ1^* mutant cilia (E’) (arrows) but the region is less well organised, and there are gaps between the ciliary and sheath cell membranes (*white* arrowheads). (**E,E’**) Electron tomograms of the area just below the ciliary BB in WT (D) and *Mks1^Δ1^* mutants (D’) reveal the rootletin filaments (arrows) that originated from the BB to surround the daughter centriole (*red* capped lines) and extend into the neuronal cell body. Several vesicles (*blue* asterisks) are visible at the base of the BB and the daughter centriole in the space delimited by the rootlelin. Serial reconstructions of this area reveal no obvious differences between WT and and *Mks1^Δ1^* mutants. (**F-G’**) Electron tomograms (F,F’) and the corresponding 3D models (G/G’) of WT (F,G) and *Mks1^Δ1^* (F’,G’) sensory cilia at 72h APF. The number of axonemal MTs has increased from 48h to 72h and mutant cilia appear more similar to WT, although an internal membranous structure can be discerned (*gold*) that is more extensive in *Mks1^Δ1^* mutant cilia. (**H,H’**) At 72h APF electron dense bridges (*black* arrowheads) connecting the MTs (*green* arrowheads) and the ciliary inner membrane are visible in both WT (H) and *Mks1^Δ1^* mutants (H’) towards the cilium tip where the electron dense crystalline-like structure is present (*blue* capped line) and appears similarly organised in both WT and *Mks1^Δ1^* mutants. Scale bars=500 nm in A, A’, B, B’, F, F’, G and G’, and 100nm in all other images.

*Mks1^Δ1^* mutant cilia (n=3) were generally thinner than WT cilia and the ciliary MTs were much sparser (Figure 5A’, B’, Movie S5). This was because most BB MTs terminated just above the TZ with only ~1-3 MTs extending into the cilium proper, compared to the 17-18 MTs observed in this region of WT axonemes (Figure 5A’, B;’ Figure 6A). The crystalline-like structure at the tip of the axoneme was also more disorganised and the connection between the crystalline structure and the ciliary/sheath membranes contained several gaps (arrows, Figure 5E’). The TZ was also noticeably shorter (Figure 5A’, B;’ Figure 6B) and the electron dense region of the TZ membrane was more diffuse than in WT (*blue* bar, Figure 5C’). The arrangement of the centriole pair and ciliary rootlets at the base of the cilium, however, appeared largely unperturbed (Figure 5D’). Thus, the ultrastructural organization of the 48h APF cilia is strongly disrupted in *Mks1^Δ1^* mutant sensory neurons.

**Figure 6.**
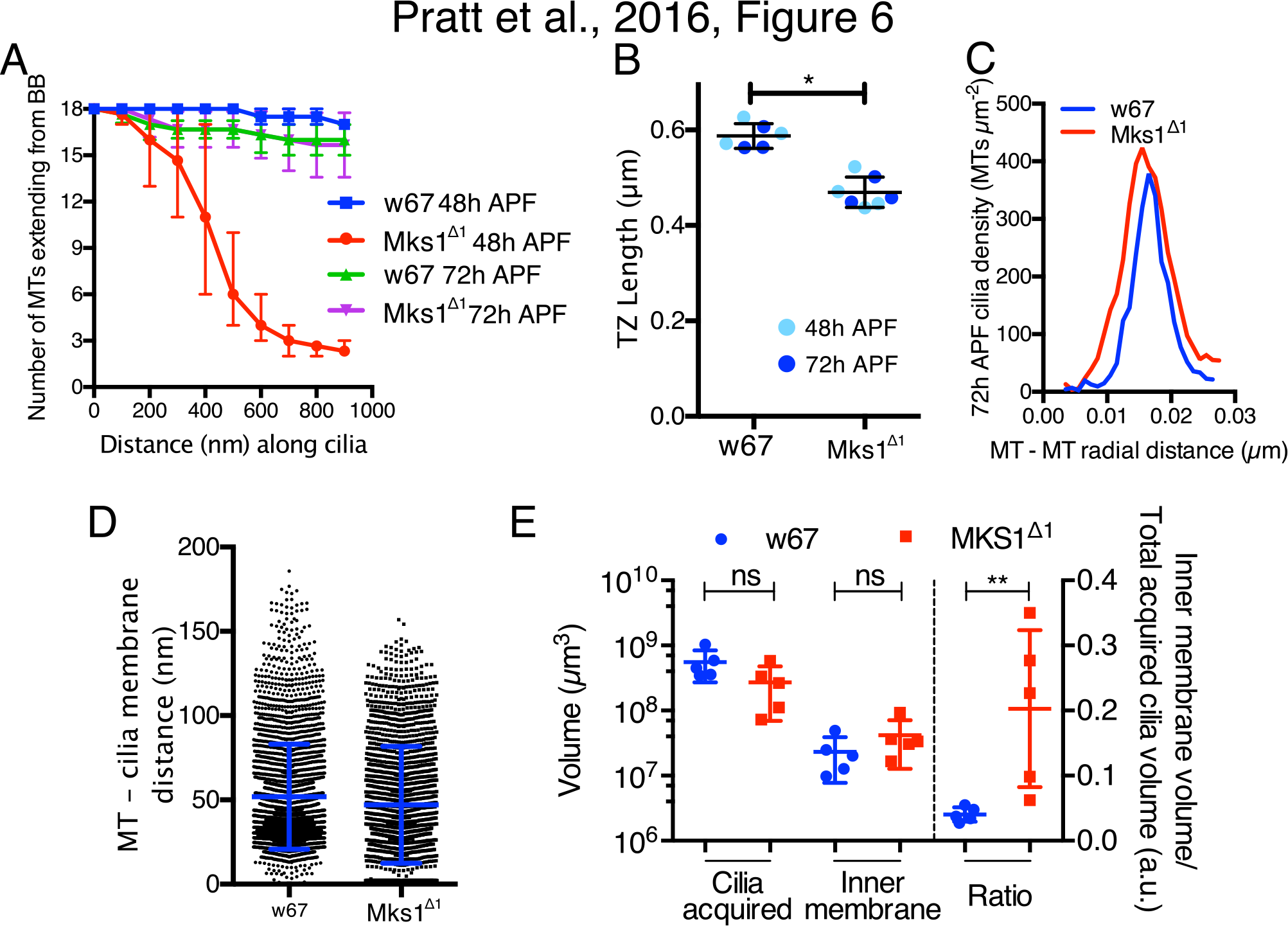
A comparison of some of the features of WT and *Mks1^Δ1^* sensory cilia. (**A**) Graph shows the quantification of the number of microtubules originating from the BB that elongate into the cilium in WT and *Mks1^Δ1^* sensory cilia at 48h and 72h APF. These data are obtained from a single cilium of each genotype, but a similar pattern was observed in multiple cilia for each genotype. (**B**) Graph shows the quantification of TZ length (measured from EM tomograms) in WT and *Mks1^Δ1^* sensory cilia at 48h and 72h APF. (**C**) Graph shows that average spacing between the ciliary MTs is ~16nm in both WT and *Mks1^Δ1^* mutant cilia at 72h APF. (D) Graph shows the quantification of distance between the cilia MTs and the ciliary membrane in WT and *Mks1^Δ1^* sensory cilia at 72h APF. (**E**). The graph shows the quantification of the total ciliary volume acquired, the volume of the cilia internal membrane acquired and the ratio of the two (all data calculated from EM tomograms of 72h APF cilia). The *Mks1^Δ1^* mutant cilia appear slightly smaller and have a slightly larger volume of internal membranes, but these differences are not significant (perhaps due to the low n-numbers as we could only reconstruct a relatively small number of cilia); the ratio of these parameters, however, reveals a clear difference between the mutant and WT. Error bars represent range in A, and s.d. in all other graphs; significance was assessed with a two-tailed t-test: ^*^p<0.01, ^**^p<0.001.

At 72h APF the overall organisation of the WT cilium (Figure 5E,F, Movie S6) was similar to that seen at 48h APF, although the volume of the cilium had increased and the cilium contained many more MTs that ran along the main axis of the cilium but that were not directly connected to the BB. Most of these MTs were in close contact with the ciliary membrane and small links between the MTs and the membranes were visible (*black* arrows, Figure 5H), similar to the membrane-microtubules connectors previously observed in chordotonal cilia (Young, 1973). These MTs had a radial separation of ~16nm (Figure 6C) and were on average ~51nm (median of ~41nm) from the ciliary membrane (Figure 6D). In addition, the amount of internal membrane present within the 72h APF cilium also increased (*gold,* Figure 5F,G; Figure 6E), and this extra membrane seemed to originate from invaginations of the neuron cell membrane into the cilium (arrow, Figure S4). The inner volume of this membrane was therefore in contact with the extracellular cytoplasm of the support cells. The membrane at these regions was very electron dense, suggesting the presence of high levels of protein material and/or saturated lipids (Figure 5E).

Interestingly, the 72h APF *Mks1^Δ1^* mutant cilia had a MT organization that was much more similar to WT than seen at 48h APF, and many of the BB MTs now extended to the tip of the cilium and large numbers of non-ciliary MTs were also present aligned along the long axis of the cilium (Figure 5,G, Movie S7). We noticed, however, that there was a larger volume of inner membrane within Mks1^Δ1^ cilia compared to WT cilia (*gold,* Figure 5F,G; Figure 6E), and the ratio of cilium volume to inner membrane volume appeared significantly increased in Mks1^Δ1^ mutants.

## Discussion

The TZ is essential for proper cilia function (Czarnecki and Shah, 2012; Reiter et al., 2012), and recent studies have suggested that an MKS-and an NPHP-module of proteins work partly redundantly with a Cep290-module to establish the TZ (Basiri et al., 2014; Schouteden et al., 2015; Williams et al., 2008, 2011). The NPHP-module seems to have arisen later in evolution and appears to be absent in *Drosophila,* but 6 potential members of the MKS-module have been identified in flies (Barker et al., 2014; Basiri et al., 2014). We show here that mutations that disrupt the function of either of two members of the MKS-module, MKS1 or B9D1, strongly disrupt the TZ-localisation of the other MKS-module components, but do not detectably disrupt the TZ-localisation of Cep290 or the TZ component Cby, strongly supporting the idea that the MKS-module proteins form a functional unit in flies. Despite the apparent lack of a functional MKS-module in *Mks1^Δ1^* or *B9D1^Δ1^* mutants, these flies exhibit surprisingly subtle cilia and flagella defects.

Although cilia function is only mildly perturbed in *Mks1^Δ1^* and *B9D1^Δ1^* mutants, the defects we do observe are informative. The MKS-module proteins are all highly expressed in testes and two types of cilia are formed in this tissue: short axonemes extend from the long centrioles found in mature spermatocytes, and these centrioles will later form the BB of the sperm flagella. In *Mks1^Δ1^* mutants the spermatocyte axonemes are dramatically reduced in length, although Cep290 and Cby are both localized normally at the base of the axoneme. The function of these spermatocyte cilia are unknown, but our findings demonstrate that the axonemes of these cilia can be dramatically shortened without any obvious effect on the fly. In particular, the subsequent formation of the sperm flagella is not detectably perturbed and mutant males appear to exhibit normal levels of fertility. This is in contrast to *Cep290* mutants that also have short spermatocyte cilia (that actually lack an axoneme), and that subsequently exhibit dramatic defects in the sperm flagella axoneme and are male sterile (Basiri et al., 2014). It therefore remains unclear why MKS-module proteins are so highly expressed in testes. We suspect that these proteins must contribute to cilia and flagella function in the testes, but that this function is not apparent under laboratory conditions in the assays we have used here. Nevertheless, it is clear the Cep290 can organize a sufficiently functional TZ in fly testes in the apparent absence of the MKS-module, but MKS-module proteins cannot do the same in the absence of Cep290 (Basiri et al., 2014).

*Cep290* mutant flies are also severely uncoordinated due to defects in their sensory cilia (Basiri et al., 2014), while we find that *Mks1^Δ1^* mutants are not noticeably uncoordinated and perform at least as well as WT flies in various assays that assess cilia function. Indeed, mutant flies appear to climb slightly faster than WT flies, and the mechano-sensory cilia that attach to the bristles on the notum are, if anything, slightly more sensitive than WT. While the distribution of the cilia membrane ion channel NompC protein was not detectably perturbed in *Mks1^Δ1^* mutant sensory cilia, the ratio of inner membrane volume to total cilium volume appeared increased in 72h APF mutant sensory cilia. These internal membranes were formed from invaginations of the plasma membrane, so the increase in membrane volume could lead to an increase in ion flow upon membrane channel opening. It is unclear, however, how this increase in inner membrane comes about. One possibility is that the lack of ciliary MTs and the looser organisation of the ciliary membrane we observe at earlier stages of development underlies the eventual enlargement of the inner membrane structures in *Mks1^Δ1^* cilia.

Notably, the localization and distribution of the IFT components RempA and NompB were also subtly perturbed in *Mks1^Δ1^* mutant cilia, indicating a potential defect in IFT—as has been suggested previously for MKS-module mutants (Zhao and Malicki, 2011). Moreover, in WT sensory cilia our EM analysis identified extended electron dense “particles” on the inner lumen of the BB that appeared to be similar to the electron dense “trains” of IFT particles previously described in axonemes and that localize between the axonemal MTs and the plasma membrane. In *Mks1^Δ1^* mutants these particles seemed to accumulate in the lumen of the BB. Clearly more work will be required to determine the identity and composition of these structures, but these observations raise the intriguing possibility that IFT particles can assemble on the inner lumen of the BB wall in these cilia. Supporting this idea, this accumulation is in agreement with our results showing that NompB-GFP localization is extended in *Mks1^Δ1^,* and with previous work that also show an accumulation of NompB in RempA mutants in the chordotonal organ cilia (Lee et al., 2008b). Together, these results suggest that *Mks1* initially affects the localization of RempA, which leads to the abnormal accumulation of NompB.

Finally, although sensory cilia structure was only moderately perturbed in *Mks1^Δ1^* mutant pupae at 72h APF, cilia structure was much more profoundly disrupted earlier in development at 48h APF. Most strikingly, the axoneme in these developing pupae had severely disrupted MTs, with very few of the BB MTs extending into the axoneme. Interestingly, it has been recently reported that an IFT B component can regulate ciliary MT growth by negatively regulating a microtubule associated protein, MAP4, that acts as a MT stabilizer (Bizet et al., 2015). Since *Mks1^Δ1^* appears to lead to an accumulation of NompB, the IFT B component in flies, a similar process could be occurring here. More work will be necessary to explain the mechanistic basis of this phenotype, but this observation strongly suggests that a properly functioning TZ is required for the efficient establishment of the axoneme. It is therefore plausible, that the MKS-module has important functions in establishing proper cilia structure and function in flies but that, given enough developmental time, cilia can compensate for the lack of these proteins and a largely functional TZ can eventually be established and maintained in the absence of these proteins. Again, we note that this is not the case when proteins like Cep290, Cby and Dila are disrupted, suggesting that the MKS1-module is functionally subordinate to these proteins in flies (Basiri et al., 2014; Enjolras et al., 2012).

Our observations suggest that flies will be an excellent system with which to probe the basic cellular functions of the MKS-module proteins. The absence of these proteins produces quantifiable phenotypes, but does not dramatically disrupt the development or viability of the organism. Such studies may shed important light on the precise roles of these proteins and how this relates to the generation of cilia pathologies in humans.

## Materials and methods

### Fly stocks

*w^67^* was used as a control in all experiments. *Mks1^Δ1^* (this study) was generated by imprecise excision of a p-element (Rubin and Spradling, 1982) 581 bp upstream of *Drosophila* Mks1, p{Epgy2}CG2556EY2042 (Bloomington). *B9D1^Δ1^* (this study) was generated by CRISPR Cas9. Three guide RNAs (gRNAs: GAAAGGTGGCCGAGACTATTTGG, CATCGTGGGCCAAATAGTCTCGG and CATCAGTCTCCCGGGCAACGAGG) were cloned into the pCFD3-dU6:3gRNA plasmid (Port et al., 2014) and injected into Drosophila embryos expressing the Cas9 gene under the control of the Nanos promoter (Port et al., 2014). *Plp^Rec5^* was published previously (Martinez-Campos et al., 2004). Mks1-GFP, B9D1-GFP, B9D2-GFP, TMEM216-GFP, Tectonic-GFP (all this study) are all expressed from the Ubiquitin promoter, which drives moderate expression in all tissues (Lee et al., 1988). CC2D2A-GFP (this study) is driven by the endogenous promoter by including a ~2 kb upstream region in the transgene. Transgenes were cloned into pUbq Gateway (Invitrogen) vectors as described before (Basto et al., 2006) labelled with *w*^+^. Transgenic lines were generated by BestGene Inc or the Department of Genetics Fly Facility (University of Cambridge). The following transgenic lines were published previously and were kind gifts: Cby-GFP (B. Durand, Enjolras et al., 2012), Cep290-GFP (T. Avidor-Reiss, Basiri et al., 2014), NompA-GFP (R. Stanewsky, Chung et al., 2001), NompB-GFP (Maurice Kernan, Han et al., 2003), NompC-GFP (Y. Nung, Yan et al., 2013), NompC-Gal 4 (Y. Nung, Yan et al., 2013), RempA-YFP (A. Jarman, Lee et al., 2008).

### Behaviour assays

To quantify male fertility, single *Mks1^Δ1^* or *w^67^* flies were crossed to 3 *w^67^* females. Hatched embryos were counted every morning and evening for 6 days. Each of the 25 biological replicates is presented as the average number of hatched embryos per day. The standard climbing assay was adapted from (Ma and Jarman, 2011). Briefly, 15 1-3 day old adult flies were knocked to the bottom of a cylinder by tapping. After 10 seconds, the number of flies above 12 cm was counted. Data is presented as the averages of 3 technical replicates. To measure the distance climbed by individual flies, 10 1-3 day old adult flies were filmed being knocked to the bottom and crawling up the sides of the cylinder. The experimental time was reduced to 7 seconds to prevent flies reaching the top of the cylinder within the experimental time frame. The distance climbed by each fly was measured in Fiji (Schindelin et al., 2012). Data is represented as the average of 5 technical replicates. For the grooming assay, 10 1-3 day old flies for each time point were covered in Reactive Yellow 86 dust (Organic Dyestuffs Corporation) by shaking them in a vial (Seeds et al., 2014). Excess dust was removed by gently tapping the flies against the mesh of a cage. The flies were then left to groom themselves for 30 or 90 minutes before taking pictures of the head and the notum on a dissecting microscope equipped with a Digital Sight camera (Nikon). The head or notum was selected as an ROI using the polygon tool in Fiji, and the area was measured. Next, the colour threshold was set to the Reactive Yellow 86 dust to select the areas covered in the dust. The area of these parts within the original ROI was also measured and the ratio was determined. Data is represented as the average over 10 flies.

### Electrophysiology

Bristle recordings were collected from individual 2-4 day old males as described in (Kernan, 1994). Humeral and notopleural bristles were cut along their midpoint and a tungsten wire reference electrode was inserted into the thorax. A glass capillary electrode containing 121 mM K^+^, 9 mM Na^+^, 0.5 mM Ca^2+^, 4 mM Mg^2+^, 35 mM glucose, and 5 mM HEPES, pH 7.1, was then placed over the cut bristle to record mechanoreceptor potentials (MRPs). MRPs were evoked by 30 μm deflections of the electrode and bristle that were generated by software-controlled movement of a PatchStar micromanipulator (Scientifica). Analog signals were acquired through a MultiClamp-700B amplifier and digitised with an Axon Digidata 1550A A/D board. Data were collected in Clampfit 10.5 (Molecular Devices) and MRP amplitudes were measured offline. Mean amplitudes from each genotype were then compared using a one-way ANOVA with Tukey′s multiple comparisons test in Prism 6 (Graphpad).

### Fluorescence microscopy

3D SIM of testis was performed as described previously (Roque et al., 2012). Live imaging was performed in Nikon Eclipse TE200-E spinning disk confocal system, equipped with an EM-CCD Andor iXon+ camera, controlled by the Andor IQ2 software. Pupae were prepared for imaging as previously described in (Jauffred & Bellaiche, 2012).

### Image analysis

The diameter of GFP tagged transition zone proteins that formed a barrel were calculated by fitting a double Gaussian to an intensity profile line perpendicular to the longitudinal axis of the axoneme, and measuring the distance between peaks. The same fitting was used to calculated the Full Width Half Maximum (FWHM) value for each peak and the average was used used as the thickness of the tube wall. For GFP tagged transition zone proteins that formed a rod, a single Gaussian curved was fitted and the FWHM calculated. Profile curves were plotted with Fiji (Schindelin et al., 2012) and fitting done with Prism 6 software (Graphpad). Distributions were tested for Gaussian distributions by the D′Agostino & Pearson omnibus test. Significance between distributions was tested by an unpaired t test for Gaussian distributions and the Mann-Whitney test for non-Gaussian distribution. Composed images of electron tomograms and the localisation maps of transition zone proteins obtained by 3D-SIM were created by scaling the Asl staining quantifications to the EM micrograph BB wall and applying the same scaling to the remaining quantifications. The intensities of NompA-GFP, NompB-GFP, NompC-GFP and RempA-GFP were all measured in Fiji by creating a sum projection, drawing an ROI around the signal and measuring the mean grey value. The background was measured adjacent to the signal in the same way and deducted from the fluorescence signal. The length of the NompB-GFP signal was measured in IMOD (Kremer et al., 1996). The signal was traced using open contours and the length of the trace was measured.

### Electron tomography

Testis samples were prepared as previously reported (Roque et al., 2012). Pupae samples for ET were prepared accordingly to a modified protocol from the testes samples. In brief, pupae were removed from their case at 48 or 72 hours APF as indicated and placed into 2.5% glutaraldehyde and 4% paraformaldehyde in 0.2 M PIPES buffer (fixative solution) and heptane for 1-2 hours rotating at room temperature. After this the pupae were transfer to 0.2 M PIPES and dissected with a pair of tungsten needles. The abdomen, head and cuticle were removed. The dissected samples were placed in fixative solution at 4°C overnight. Samples were washed 3x for 15-30 minutes in 0.2 M PIPES. En-bloc staining was performed in 1% OsO_4_ in miliQ H_2_O for 90 minutes at room temperature. This was followed by 5 washes in miliQ H_2_O for 5 minutes each. Secondary fixation was performed in 1% uranyl acetate solution in milliQ water overnight. This was followed by 3 washes in miliQ H2O. Dehydration was performed in a series of 30%, 50%, 75% and 100% EtOH for 45-60m each, followed by a step in 100% acetate for 45 minutes. Embedding was performed in a series of 25% (2 hours), 50% (2 hours) and 75% Agar100 in acetone (overnight), and 100% acetone overnight. After a fresh 100% Agar100 change the samples were polymerised at 60°C for 24 hours. Tilt-series were acquired with SerialEM (Mastronarde, 2005) in a Tecnai T20 at 120Kv and tomograms reconstructed and modelled with iMOD (Kremer et al., 1996).

## Acknowledgements

We thank members of the Raff laboratory for advice and comments on the manuscript, the Micron Oxford Advanced Bioimaging Unit (supported by a Wellcome Trust Strategic Award (091911) for access to the OMX 3D-SIM system, and Errin Johnson for EM assistance at the Dunn School Bioimaging Facility. M.B.P. was supported by a BBSRC PhD Studentship, J.W.R and H.R. were supported by a Wellcome Trust Senior Investigator Award (104575), J.T. and I.D were supported by a Wellcome Trust Senior Research Fellowship (096144); A.R.B. and H.R.D. were supported by a grant from the Medical Research Council (G1001644).

M.B.P. and H.R. designed and performed the majority of experiments and helped to write the manuscript. I.D. and J.T. designed the electrophysiology experiments and J.T. performed the electrophysiology experiments. A.R.B. and H.R.D. devised and performed the bioinformatics analysis that identified the genes studied here. J.W.R. helped design experiments and helped write the manuscript.

## Supplementary Figures

**Figure S1.**
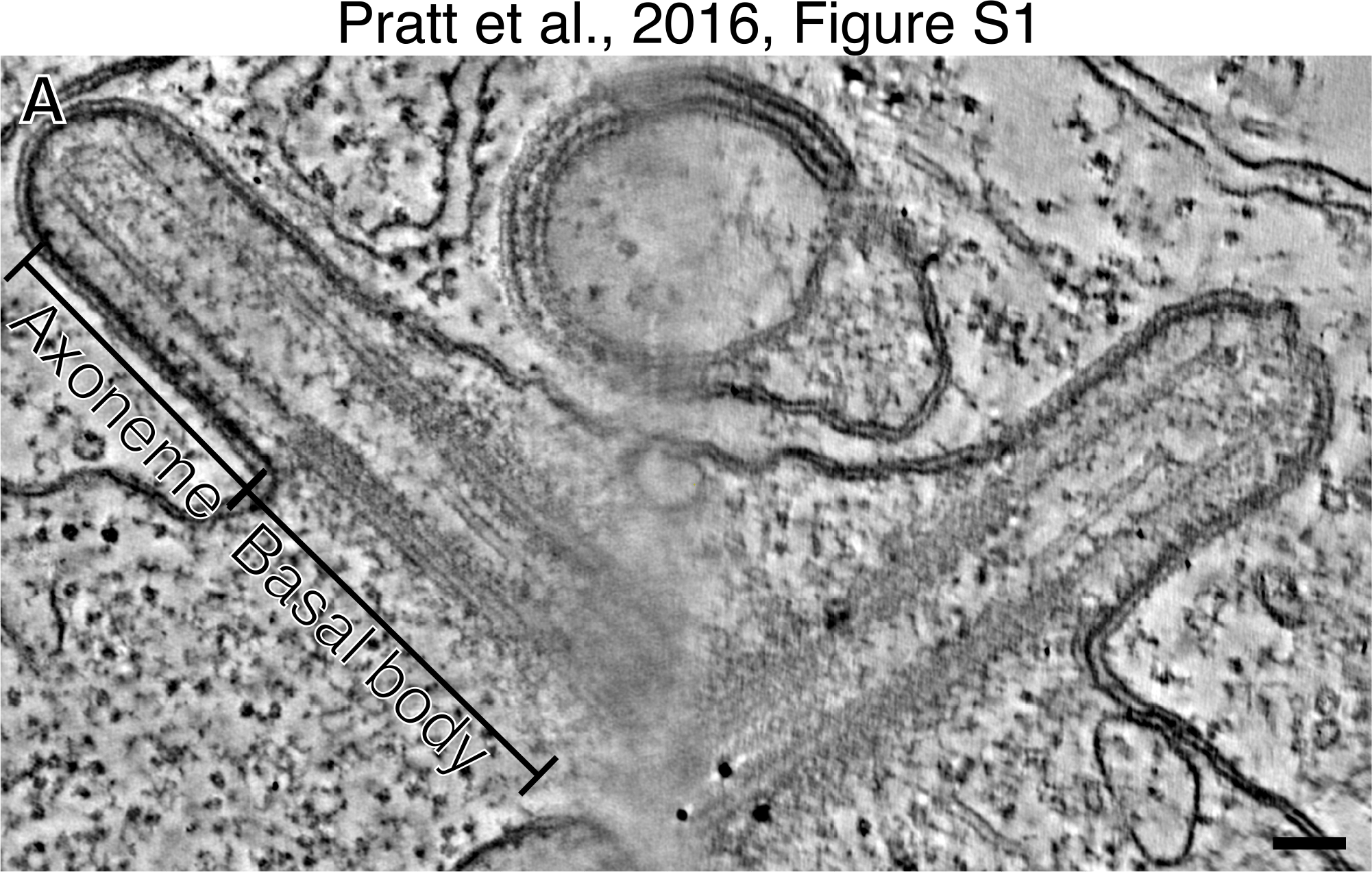
An EM tomogram of a WT spermatocyte axoneme. Both BBs extend a characteristic short axoneme in this cell type.

**Figure S2.**
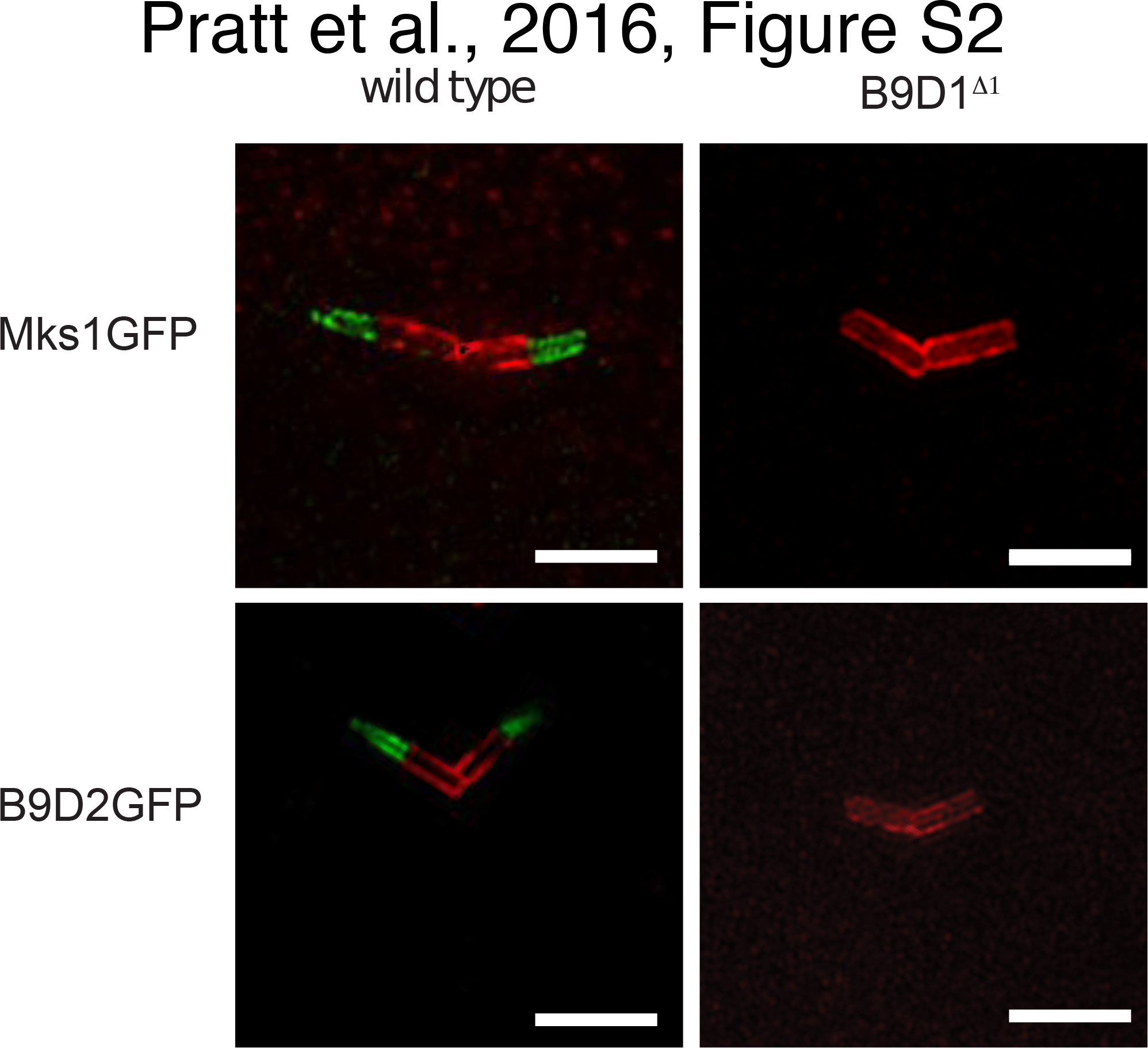
MKS-module proteins are not detectable at the TZ in *B9D1^Δ1^* axoneme. GFP tagged MKS1 and B9D2 localise to the axoneme in wild type (left panel) spermatocyte axonemes, but are not detectable in *B9D1^Δ1^* mutant axonemes (right panel).

**Figure S3.**
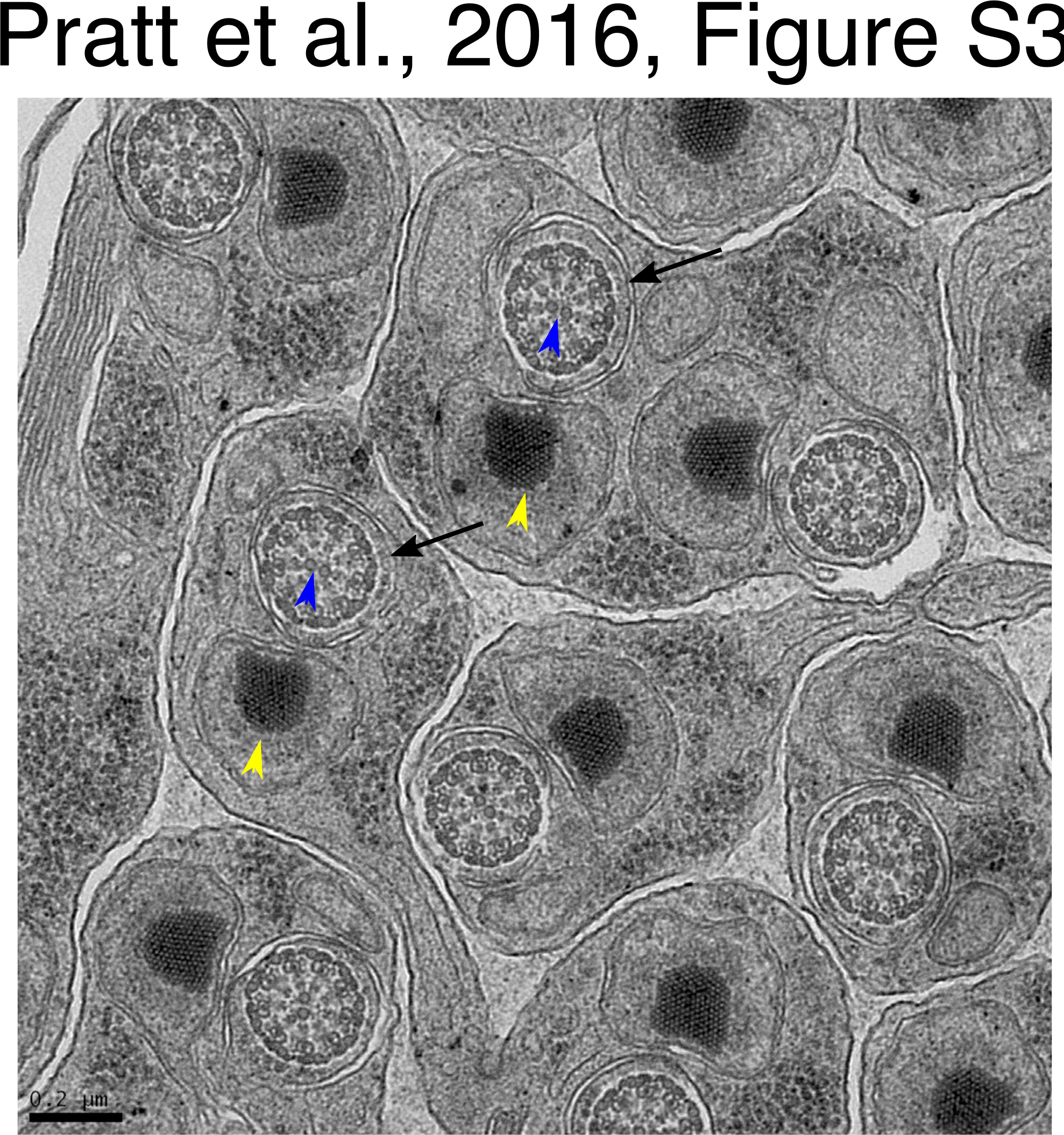
Mks1^Δ1^ flagella are structurally normal. Flagella from Msk1^Δ1^ mutant have a normal nine-fold symmetry (arrows), with the clearly central pair visible (blue arrowhead) as well as the mitochondria body (yellow arrowhead).

**Figure S4.**
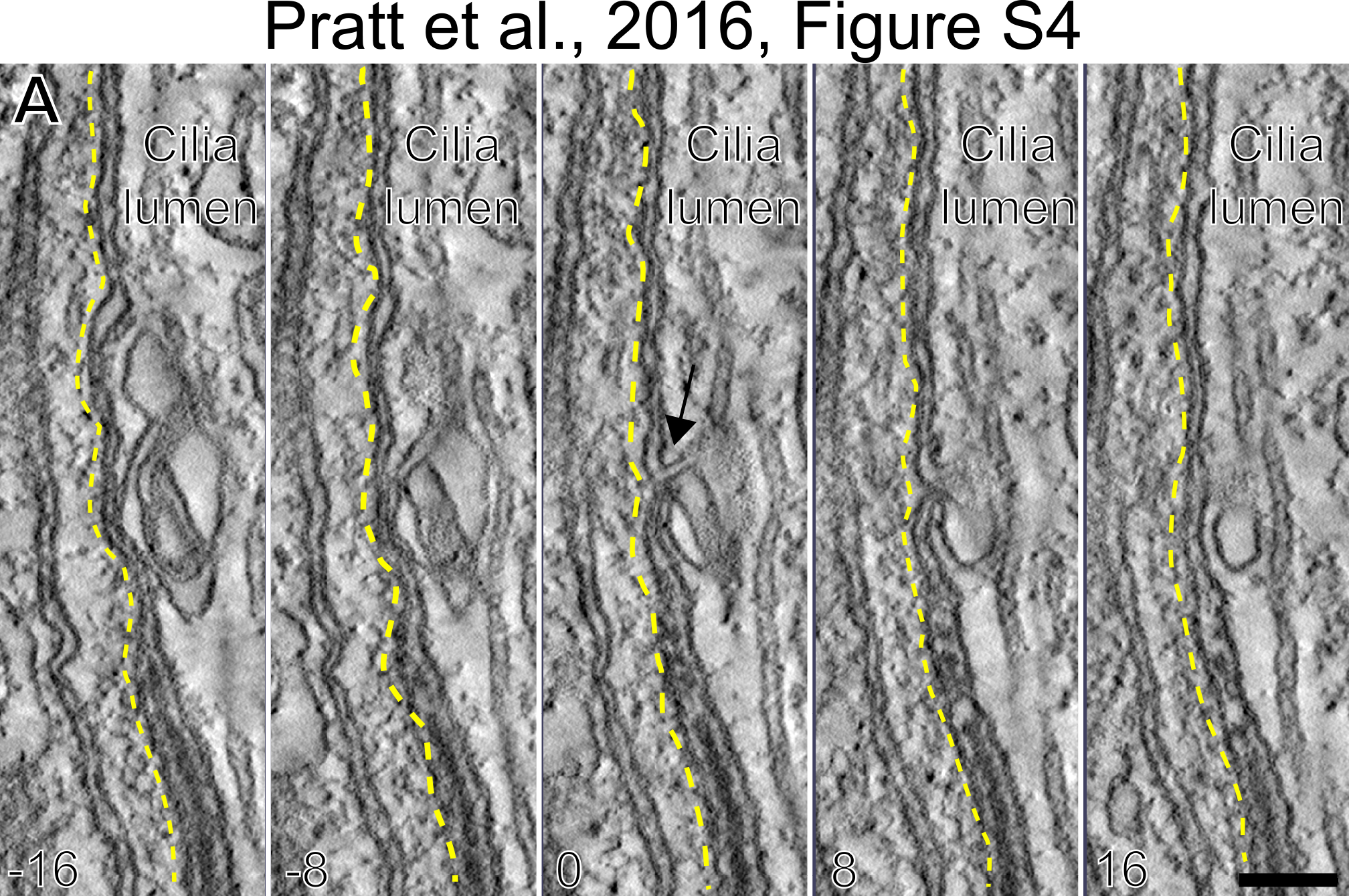
The inner ciliary membrane appears to be contiguous with the outer plasma membrane of the sensory neuron. Micrographs show serial sections of the same electron tomogram from a 72h APF sensory cilia. The convoluted membrane inside the cilia can be visualised (arrowheads), and this membrane can be observed to fuse directly with the ciliary membrane (arrow), which is directly abutted to the sheath cell membrane (yellow dotted lines). The micrographs were taken every 8 sections before and after the middle panel (bottom left numbers). Scale bar=100nm.

## Movie S1

Electron tomogram of wild type cilium of primary spermatocyte, corresponding to Figure 3A. Scale bar=100nm.

## Movie S2

Electron tomogram of Mks1^Δ1^ cilium of primary spermatocyte, corresponding to Figure 3A’. Scale bar=100nm.

## Movie S3

Electron tomogram of Mks1^Δ1^ Mks1-GFP rescue of cilium of primary spermatocyte, corresponding to Figure 3A’. Scale bar=100nm.

## Movie S4

Electron tomogram and corresponding model of a wild type sensory cilium of a bristle notum, corresponding to Figure 5A, B. Scale bar=250nm.

## Movie S5

Electron tomogram and corresponding model of a Mks1^Δ1^ mutant sensory cilium of a bristle notum, corresponding to Figure 5A’, B’. Scale bar=250nm.

## Movie S6

Electron tomogram and corresponding model of a wild type sensory cilium of a bristle notum, corresponding to Figure 5F, G. Scale bar=250nm.

## Movie S7

Electron tomogram and corresponding model of a Mks1^Δ1^ mutant sensory cilium of a bristle notum, corresponding to Figure 5F, G. Scale bar=250nm.

